# Evidence of sheeppox in 17^th^-19^th^ century France: a multi-disciplinary investigation of a sheep mass mortality assemblage

**DOI:** 10.1101/2025.11.24.690131

**Authors:** Annelise Binois-Roman, Louis L’Hôte, Roisin Ferguson, Kevin G. Daly

## Abstract

Epizootic outbreaks posed major threats to food security, economic stability, and animal welfare in past communities. While these events and their impacts are well documented in historical records, very few archaeological examples are known, and none for which the causal agent has been reliably identified. We therefore investigated a 17 ^th^–19^th^ century ovine mass-mortality assemblage from Louvres, France, using an interdisciplinary approach integrating paleogenetics, zooarchaeology, history, and veterinary science.

The deposit contained nine complete, articulated sheep skeletons, mostly older individuals of both sexes, which were unbutchered but displayed flaying marks. Aside from minor age-related lesions and one ectopic bone nodule, the animals exhibited good skeletal health, with no obvious cause of death. Ancient DNA analysis was therefore conducted and proved highly successful, conclusively identifying two ovine pathogens: sheeppox virus (SPPV) and the parasite *Taenia hydatigena*. When considered alongside archaeological, veterinary and historical evidence, these findings indicate that the animals died during a sheeppox outbreak.

This study provides the first detection of SPPV in archaeological bone and the first paleogenome of this virus. It also offers new insights into livestock health, mortality, and disease management in early modern France, demonstrating the value of integrated approaches for reconstructing past epizootics.

## Introduction

“*Thicker than squalls swept by a hurricane from off the sea, plagues sweep through flocks; and not one by one diseases pick them off, but at a stroke, a summer’s fold, present and future hopes, the whole stock, root and branch.”* (Virgil, Georgics III, ll.471-473,[1]) Written by Virgil in the first century BCE, these verses highlight well the major concern that epizootic outbreaks represented for ancient sheep farmers. And rightly so: many depended on the flock’s production - including wool, cheese, meat - for their subsistence, and the loss of their animals would have had potentially devastating economic and sanitary consequences. Animal disease can still have similar impacts on societies today, an issue which the One Health framework developed at the turn of the millennium has brought into focus in recent years [2]. But although the effect of animal disease on present day societies - whether through direct zoonotic transmission or indirect economic and social repercussions - has been studied in depth, our understanding of its impact on past societies, which relied more heavily on animal resources and lacked our modern medical arsenal, remains fairly limited.

Textual evidence of epizootic animal mortalities is encountered in almost every written civilisation, and invariably highlights the severity of these events [3–5]. Material and skeletal evidence is, on the other hand, few and far between, its study plagued by technical and methodological issues. The disarticulated, butchered state in which animal remains are typically found is an important limiting factor, but even mass animal burials are often difficult to interpret, requiring to distinguish between disease, ritual depositions and accidental mortalities [6,7]. Recent advances in paleogenetics have facilitated the previously near-impossible task of identifying pathogenic organisms in animal remains, with analyses now sensitive enough to detect traces of pathogen genomes both bacterial and viral e.g. [8–10]. However, demonstrating infection by an agent does not necessarily translate into the host having died from that specific cause, compounding the difficulty of the issue.

We nonetheless suggest that, given the right circumstances, archaeological epizootic mortalities can be reliably identified by an interdisciplinary approach relying on archaeological, osteological, paleogenetic, historical and veterinary lines of evidence. This paper presents how this systematic approach was applied to a sheep mass burial from historical Northern France, and allowed the identification of the animals’ cause of death - specifically here, an outbreak of sheeppox. This unprecedented result offers new insights into livestock health, mortality, and disease management in 17^th^–19^th^ century France, demonstrating the potential of integrated approaches in reconstructing past epizootics, and enhancing our understanding of both past disease expression and societal responses.

## Background

### The sheep-pit from Louvres: osteological analysis

In 2012, a small-scale archaeological assessment was carried out on a farm first built in the 16 ^th^ century CE in Louvres, France, about 25 kilometres north of Paris, with five trenches dug in what used to be the farm’s vegetable garden [11] (Figure 1). A bowl-shaped pit (feature 3, ca. 1.1 m diameter x 0.4 m deep) was brought to light, and found to contain articulated animal skeletons (Figure 2). No other noteworthy features were identified by the survey, and no excavations of the plot were mandated.

**Figure 1.**
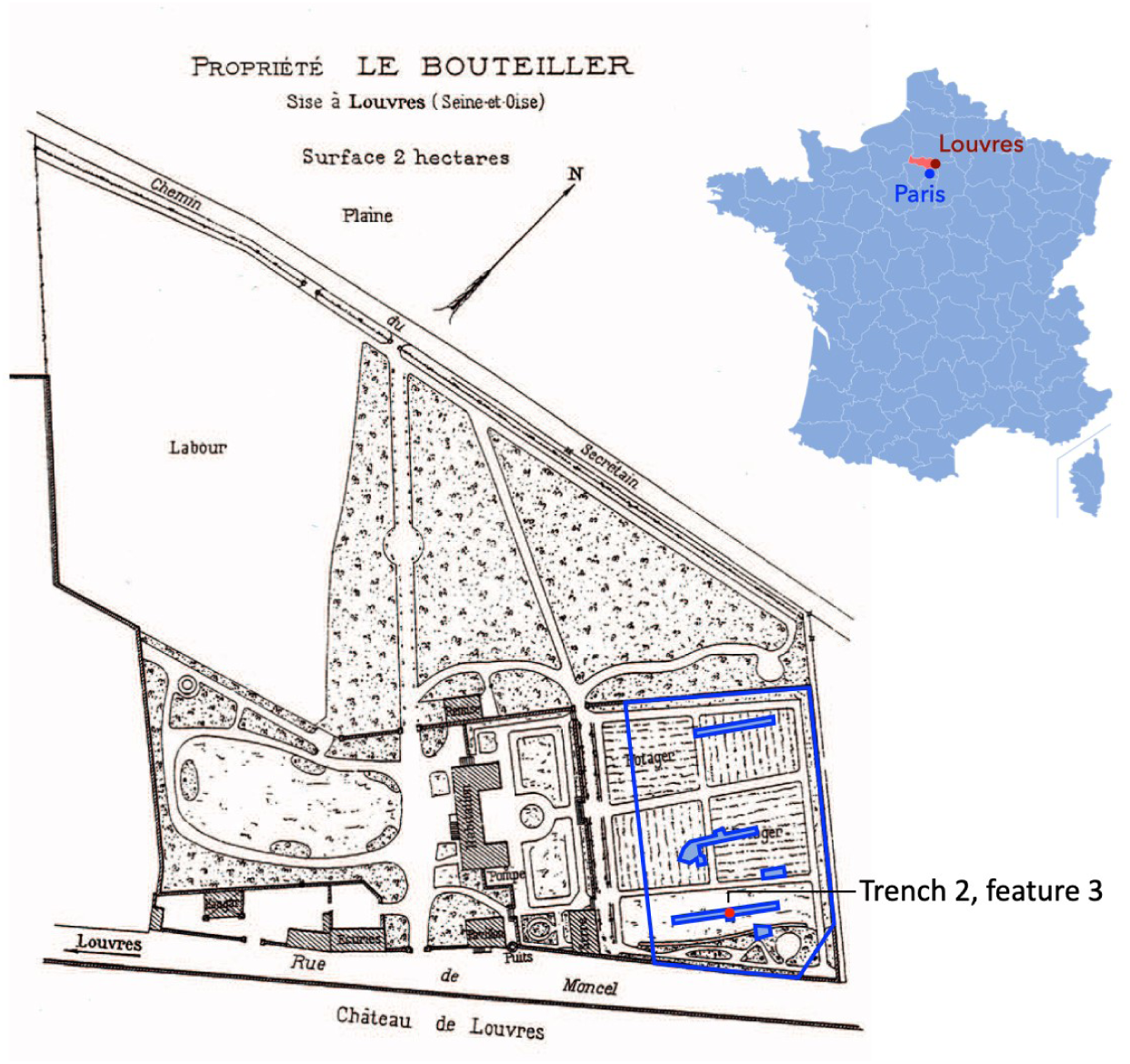
19^th^ century map of ‘le Bouteiller’ farm in Louvres, France (Bibliothèque du Musée Condé, Chantilly, in [11]). Location of the surveyed plot indicated by blue outline, the five trenches in light blue, and feature 3 by a red dot.

**Figure 2.**
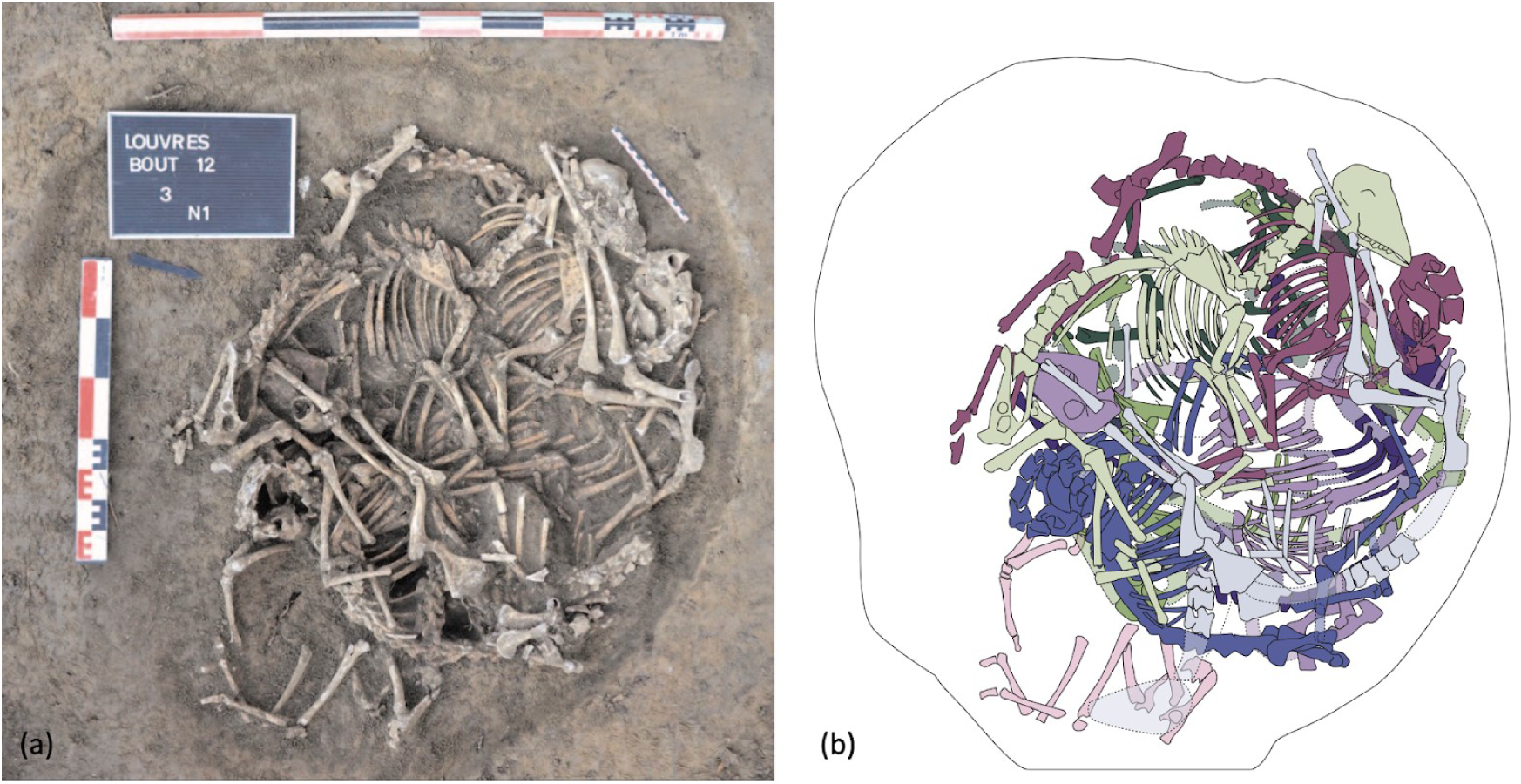
Feature 3, Louvres ‘le Bouteiller’ 2012 excavations. (a) Photograph of the excavated pit (courtesy of A. Konopka) and (b) illustration of the individual skeletons.

The skeletons were radiocarbon dated to the 17 ^th^ to 20^th^ century CE in a discontinuous probability distribution (175±30 BP, Poz-85860, calibration 2σ IntCal20 [12] in Table S1, see Figure S1). Archival evidence of land use and occupation suggests the most recent end of the distribution to be unlikely, and a probable deposit date of 1658-1888 cal CE was retained.

Careful excavation and a thorough photographic documentation of the dismantlement of the bone accumulation allowed the identification of nine skeletons, which were numbered top-down in reverse deposition order. The animals laid on their sides, with extended limbs and heads in various positions, and appeared to have been organised in the pit to optimise its infilling, with vertebral columns aligned on the edges and limbs oriented towards the centre in a circular pattern (Figure 2). Excavation data suggest the carcasses were deposited in a single episode and that the pit, apparently purpose-dug for the event, was immediately filled in.

The bone accumulation was dismantled partly by individual, partly collectively, and not all elements could be reattributed to specific skeletons post-excavation; individuals comprise therefore between 28 and 98 elements each, with over 250 elements (mostly vertebrae, ribs and phalanges) remaining non-attributed (Table 1). Bone preservation was mostly good, though the three lower-most skeletons presented some brittle elements with partly-dissolved surfaces and low bone density.

**Table 1.** Osteoarchaeological analysis of the sheep skeletons from feature 3, Louvres ‘le Bouteiller’.

| Individual ID | DNA ID | Minimum elements | Withers height (cm) | Sex (osteological) | Sex (DNA) | Age at death (years) | Cut marks | Pathology |
| --- | --- | --- | --- | --- | --- | --- | --- | --- |
| Ind 1 | RF002 | 72 | 63.5 | Male | Male | 4.5-6 | No | Minor |
| Ind 2 | RF003 | 94 | 54.9 | Female | Female | > 7 | Yes | Yes |
| Ind 3 | N/A | 75 | 59.4 | Female | N/A | older adult | No | Minor |
| Ind 4 | RF004 | 98 | 61.2 | Ambiguous | Female | 4.5-6 | No | Minor |
| Ind 5 | N/A | 28 | 56.4 | Female | N/A | older adult | Yes | No |
| Ind 6 | RF006 | 88 | 59.6 | Female | Female | adult | No | No |
| Ind 7 | N/A | 38 | 61.9 | Male | N/A | older adult | No | Minor |
| Ind 8 | RF005 | 42 | 63.4 | Male | Male | 2.5-3.5 | No | Minor |
| Ind 9 | RF009 | 34 | 61.9 | Male | Male | adult | No | No |

A zooarchaeological examination of the skeletons (see Methods) confirmed the animals were all domestic sheep (*Ovis aries* L. 1758) of average size and slender constitution (average metacarpal slenderness inde× 10.3%). Their estimated withers height varies between 54.9cm and 65.3cm (Table 1), consistent with recorded heights for Northern France at the time (Bouteiller average: 60.4cm; 17^th^-18^th^ c. average: 59.0cm; 18^th^-19^th^ c. average: 63.5cm; [13]). Sex was identified from osteological criteria on the pelvis then confirmed by DNA for six individuals (see Methods and Table 1); five females and four males were identified, all of which were hornless. The male individuals displayed fairly subtle masculine characteristics for their age, and it is likely all were castrated, as was traditional in 18^th^-19^th^ century French farming systems [14]. Age-at-death indicated an overall mature population, consisting of two prime adults (4.5-6 years old), four older adults (>7-8 years old), two unspecified adults (> 4.5 years old) and one subadult (2.5-3.5 years old). This demographic pattern in which a high number of older males are retained in the flock is suggestive of a farming system with a strong focus on the production of wool, as can be observed ethnographically in a few traditional systems [15].

Cut marks were observed on two skeletons. In sheep #5, both radii exhibited a deep incision on the proximal diaphysis, oriented perpendicular to the shaft. As the carcass was otherwise unbutchered and fully articulated, these marks are interpreted as traces of flaying, albeit situated unusually high on the limbs. In sheep #2, incisions were recorded on the atlas, consisting of a series of fine, parallel cuts on the underside of the atlanto-occipital joint. Given that the head and neck were found in close articulation, these traces cannot be attributed to beheading. Instead, they are most plausibly interpreted as evidence of slaughter by throat-slitting. However, such a procedure typically produces a single incision lower along the cervical column; the multiple cuts observed here indicate a repeated action unnecessary for that purpose. The slaughter marks on sheep #2 are therefore either atypical examples of this practice or the result of an alternative activity that remains to be identified.

The sheep from the pit displayed good skeletal health for animals their age. Four individuals showed no pathological modifications, and four showed only minor healed lesions (Table 1). Only sheep #2 presented any pathological element of note, a small calcified nodule that was found stored with its head and cervical column; its skeleton was otherwise healthy.

### A mass mortality event

Burial is almost never the intended outcome for livestock; whether after a brief fattening period or a lifetime of labour, most are ultimately slaughtered, butchered, and consumed. The burial of a farm animal therefore constitutes an anomaly in the expected course of events, and multiple burials—in which several individuals are interred together—even more so. Broadly, two phenomena account for the immense majority of multiple burials of domestic animals: mortality crises and ritual depositions [16].

It is therefore essential to distinguish between these two scenarios before further examining the Louvres “Bouteiller” sheep burial. As outlined by Roman-Binois [16], a mortality crisis can only be reliably identified when the following conditions are met: the victims belong to a single species; their number is such that unrelated, concurrent deaths are highly improbable; the deaths can be demonstrated to have been simultaneous; and the burial shows no evidence of ritual activity. The Louvres deposit meets all of these criteria, with nine sheep—above the suggested minimum of six—found densely packed within a single feature. The high degree of carcass entanglement, the perfect anatomical articulation even in the deepest layers, and the absence of weathering or scavenging marks indicate a single act of deposition followed by immediate backfilling. No material evidence of ritual activity was observed; modern-period Catholic France is moreover an unlikely setting for such activities. We therefore conclude that the Louvres sheep represent the victims of a mass mortality event.

If so, the remaining question is that of the reason for the sheep’s deaths. Mortality events can stem from a variety of causes: disease is of course common, but weather-related events and accidental deaths must not be discounted. In fact, historical evidence suggests that illnesses only accounted for about half of mortality events in pre-industrial flocks, with as many events arising from harsh winters, springtime storms, floods and predator attacks [16]. A three-pronged interdisciplinary approach was adopted to address this potential diversity of causes. Written records from modern-period northern France were examined to identify the most frequent types of sheep mortality and their clinical manifestations in that specific context. In parallel, paleopathological and paleo-epidemiological analyses of the skeletal remains were conducted to detect evidence of trauma or disease, and paleogenomic analyses were undertaken to search for biomolecular traces of pathogens. The results from these three lines of evidence were then integrated to support a final diagnosis of the sheep’s cause of death.

Only one pathological specimen was available for analysis, the nodule found with sheep #2. However, the absence of visible lesions on the skeletons does not preclude them delivering vital clues as to the cause of death – though most mortality crises intervene too swiftly to allow bones to react macroscopically to aggression, paleo-epidemiological and paleogenomic evidence can still be obtained from them. The diagnostic process adopted to address both types of material evidence is however slightly different, and they will therefore be presented subsequently in this paper.

## Results

### Analysis of the pathological nodule

The pathological nodule was recovered from among the cranial debris of individual #2, and was initially presumed to have originated from the cranial or cervical region of that animal (see Note 1 in Supplementary Information). It was a flattened, sub-circular biological calcification measuring approximately 1 cm in diameter and 0.4 cm in height. Its surface was beige and bone-like, smooth in texture but uneven in form, and punctured by several sub-circular perforations with rounded edges. The specimen was of very low density and appeared at least partially hollow. Although no photograph of the object is available, its external appearance closely resembled that of the biological artefact illustrated by Komar and al. [17]. Radiographs taken using a dental radiographic machine revealed a heterogeneous, poorly organised structure composed of voids and snowy areas traversed by a few denser lines (Figure 3). The radiographic density of the material matched that of bone.

**Figure 3.**
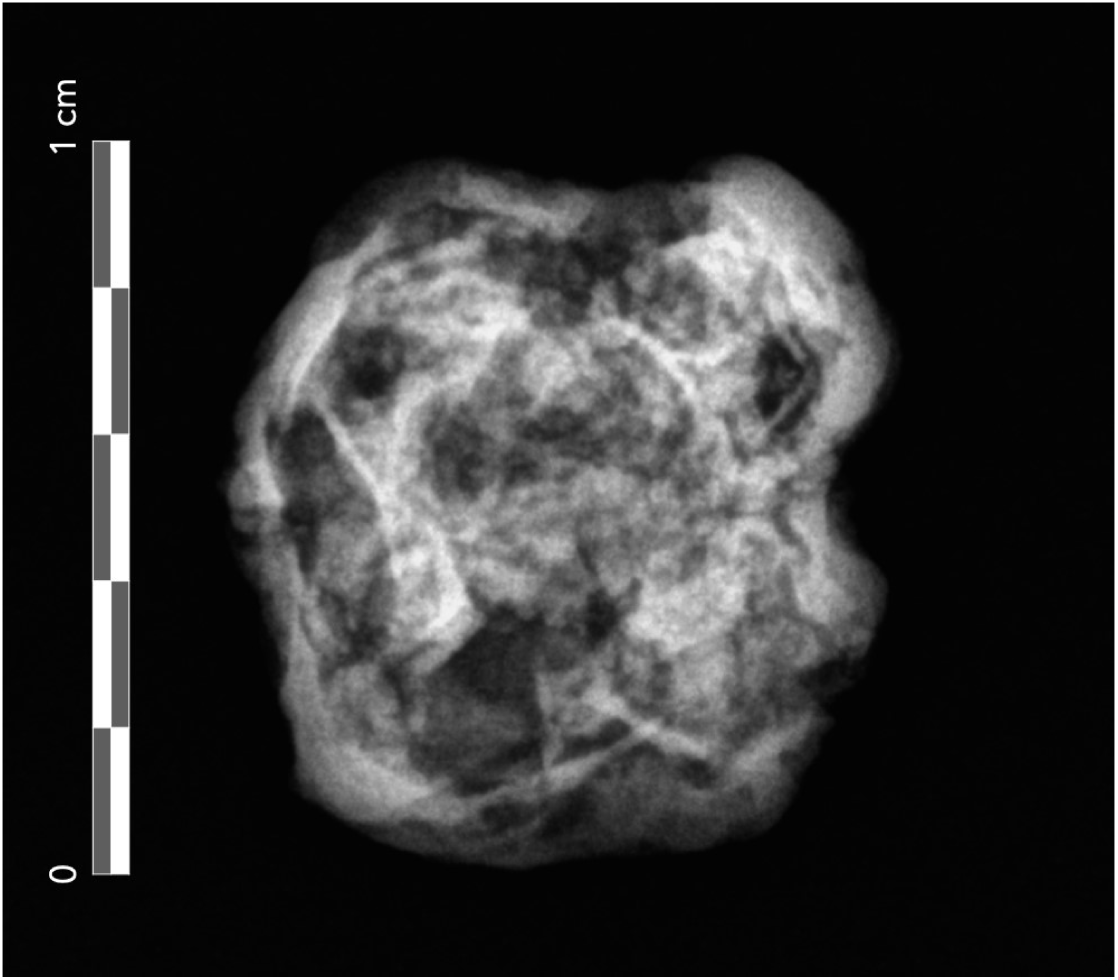
Radiograph of the nodule found with individual 2, Louvres ‘le Bouteiller’. Approximate scale overlaid onto radiograph.

The nodule’s appearance and presumed anatomical origin suggested a primary diagnostic hypothesis of a calcified cervical lymph node, which is most often in sheep caused by caseous lymphadenitis, an infectious disease due to *Corynebacterium pseudotuberculosis* (see Note 1 in Supplementary Information). A paleogenomic investigation of the specimen was therefore undertaken to clarify its origin and identify potential pathogens (see Methods). The host genome was confirmed to be sheep, with an endogenous DNA content of 28.01%, consistent with previous reports for nodule remains [18]. The individual was female and shared a mitochondrial DNA haplotype with sheep #2 (Figure S2).

To verify the genetic identity of the nodule, a D statistic test [19] was performed between sheep-aligned DNA from the nodule and four of the Louvres sheep (see below; Table S4). We posited that in tests where the nodule and the animal host of the node were included, there would be a substantial excess of “derived” genetic variants shared between these two samples. The test yielded an average absolute D statistic of 0.588 when the nodule (RF001) was placed in the H3 position and sheep #2 (RF003) was placed in the H2 position, while tests without sheep #2 ranged in absolute D values 0.0005 to 0.0294. This strongly suggests that the two samples are genetically identical and allowing secure attribution of the nodule to sheep #2. Post-mortem damage patterns at read ends confirmed the authenticity of the sheep-aligning ancient DNA (Figure S3).

Unaligned reads were examined using a unique *k*-mer counting approach [20]. Surprisingly, whilst no DNA from infectious bacteria was identified, abundant reads from a parasitic tapeworm, *Taenia hydatigena*, were detected in the sample (16,593 reads mapped, 0.0029× genome coverage). After further sequencing, alignment to the mitochondrial genome produced 75.4-fold of coverage, enabling phylogenetic confirmation of the species as *T. hydatigena* and placement within the A haplogroup (Figure 4B). DNA damage mapping across both the nuclear and mitochondrial genomes confirmed its ancient origin.

**Figure 4.**
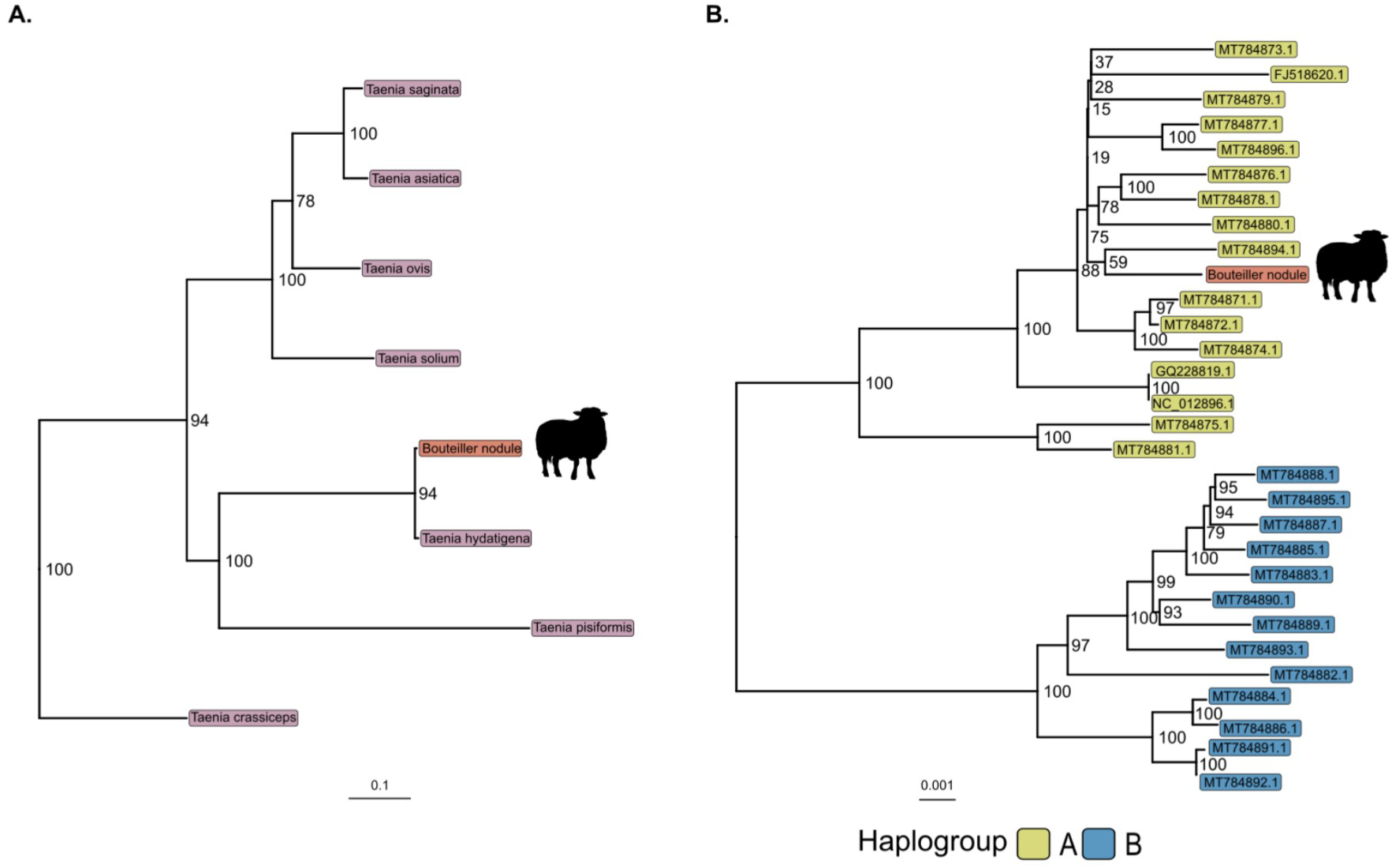
Phylogenetic placement of the Bouteiller nodule isolate among *Taenia* species and mtDNA haplogroups. **(A)** Phylogenetic placement of the Bouteiller taenid mitochondrial genome within the diversity of *Taenia* species. A maximum-likelihood phylogeny was constructed with IQ-TREE2 (21) using complete mitochondrial genomes from representative *Taenia* species. Bootstrap support values are shown at the corresponding nodes. The Bouteiller nodule sequence is highlighted in red and accompanied by a sheep silhouette. **(B)** Haplogroup assignment of the Bouteiller taenid mitochondrion. A high-resolution Maximum Likelihood mitochondrial haplogroup tree of *T. hydatigena* was generated with IQ-TREE2. Samples are coloured according to haplogroup (A= yellow, B = blue), and tip labels correspond to GenBank accession numbers (Table S7). The Bouteiller nodule sequence (red) clusters within haplogroup A. Bootstrap support values are shown at the corresponding nodes.

*T. hydatigena* is today in Europe an uncommon intestinal parasite of domestic dogs and other carnivores, having largely disappeared from industrialised countries in the 20^th^ century due to antiparasitic treatments and changes in animal-feeding practices. The parasite is acquired by the carnivorous final host through the consumption of the raw viscera of large herbivores, who serve as intermediate hosts for this parasite. As such, sheep and goats, and less commonly cattle, deer and pigs, will host the larval stage of the flatworm, previously known as *Cysticercus tenuicol/is,* on the peritoneal surface of their abdominal organs, most notably the liver, the spleen, the mesentery and the omentum (22-24). The larvae form a bladder-like cyst of 1-7 cm diameter, attached to the organs by a thin stem; in sheep, the cysts can also more rarely develop inside the liver. The disease in ruminants is chronic, and the cysts may calcify over time [25,26].

No direct prevalence data is available for *T. hydatigena* in 17^th^–19^th^-century France, but the parasite was likely widespread. It remains common where wild carnivores and stray dogs act as reservoirs and livestock are not routinely treated; reported prevalence in sheep range from 2% (India [27]) to over 50% (Tanzania [28], Ethiopia [29]), and in dogs from around 10% (Vietnam [30], Italy [31]) to over 60% (Mongolia, Dazan 1978 in [32]). Genomic evidence of the parasite has also been found in archaeological contexts, with identifications reported from a Danish Iron Age pond [33], from medieval Norse Greenland [34], and from early Modern urban refuse layers in Denmark and Lithuania [35]. As taeniid eggs cannot be distinguished to species level morphologically, no specific paleoparasitological evidence of *T. hydatigena* eggs is available, but they are likely included among the many taeniid eggs identified in archaeological deposits worldwide.

The identification of the nodule as a calcified parasitic cyst of a *T. hydatigena* larva is therefore consistent with most of our data. This parasite is a documented source of calcified cysts in sheep, and appears to have been common at the time and place of the Louvres burial. No literature regarding calcified *T. hydatigena* cyst morphology was located, probably due to its limited diagnostic relevance, but published descriptions of other taeniid calcified cysts, notably of *Echinococcus sp.*, match the Louvres specimen reasonably well. The only inconsistent diagnostic element was the presumed anatomical location. While aberrant cyst localisations have been recorded in sheep [36,37], none are in the head or cervical region, which are physiologically unsuitable for larval development. It is therefore likely the specimen was misassigned during excavation or post-excavation handling.

To our knowledge, this is the first archaeological identification of a *T. hydatigena* cyst and the first confirmed animal parasitic cyst in the archaeological record. The species is mentioned by Baker and Brothwell as a potential source of archaeological pathological specimens, but these are considered ‘most unlikely to be recovered’ [38]. Parasitic cysts have been reported in human remains—mostly interpreted as *Echinococcus* sp. infestations—at Neolithic Siberian [39], medieval Italian [40], and medieval Spanish sites [41,42], though these specimens tend to be larger and hollower than the Louvres example.

In sheep, *T. hydatigena* infestation is generally asymptomatic and usually detected incidentally at slaughter. Adult sheep may experience mild production losses, and severe infestations can occasionally cause sporadic deaths in lambs, but such events are rare [31,43]. Even so, the complaint is an individual one, never resulting in multiple deaths, especially not in mature adults. It therefore seems highly unlikely that the infestation documented in Louvres could have resulted in the death of sheep #2, let alone nine adult sheep over a short span of time. It is even more implausible that the sheep would have been culled and discarded because of a parasitic infestation. Before the late-19th century understanding of helminth lifecycles and the mid-20th century introduction of synthetic antiparasitic drugs, parasitic burdens in European livestock were extremely high. Most, if not all, animals would have harboured multiple worm species without raising concern, and infestations were rarely noted prior to slaughter. The Louvres *T. hydatigena* cyst is therefore best interpreted as an incidental finding of intercurrent disease, unrelated to the primary cause of death in the mortality assemblage.

### Analysis of the skeletal assemblage

If the nodule could not provide answers to the cause of death, perhaps the skeletons would. We first approached the diagnosis through a paleo-epidemiological angle: having identified in relevant historical sources the most common causes of sheep mass mortalities in the Modern period, we proceeded to map the epidemiological characteristics of each, including terrain, seasonality, and demography of victims (see Methods). The resulting profiles were then compared to that of the Louvres deposit in order to identify the diagnostic hypotheses that most closely matched our observations.

Unfortunately, our differential diagnosis did not lead to any conclusive results. The lack of data in Louvres relating to the season of death hindered our approach, and the atypical demographic profile, highly skewed towards older animals, contradicted most diagnostic hypotheses. Apart from the average age at death, the epidemiological profile of the deposit was fairly unspecific (Table S2); agreement was weakest with pathological causes (anthrax, liver fluke, sheeppox, red disease), and a toxic or accidental incident was initially tentatively suggested [16].

These unsatisfactory results were therefore complemented with a paleogenomic approach. A petrous bone was sampled from all available crania, and teeth presenting intact closed roots were selected from sheep #1, 2, 4, 6/7, 8 and 9 (Table 2, Table S3) and sampled within the pulpar cavity. All samples were run through an initial shotgun sequencing analysis following the protocol outlined in the Methods section; all yielded sufficient amounts of endogenous DNA (16.9% to 49.5% for the petrous, 0.02% to 20.6% for the teeth) that were aligned to sheep and confirmed to be of ancient origin through damage pattern mapping (Figure S3).

**Table 2:**
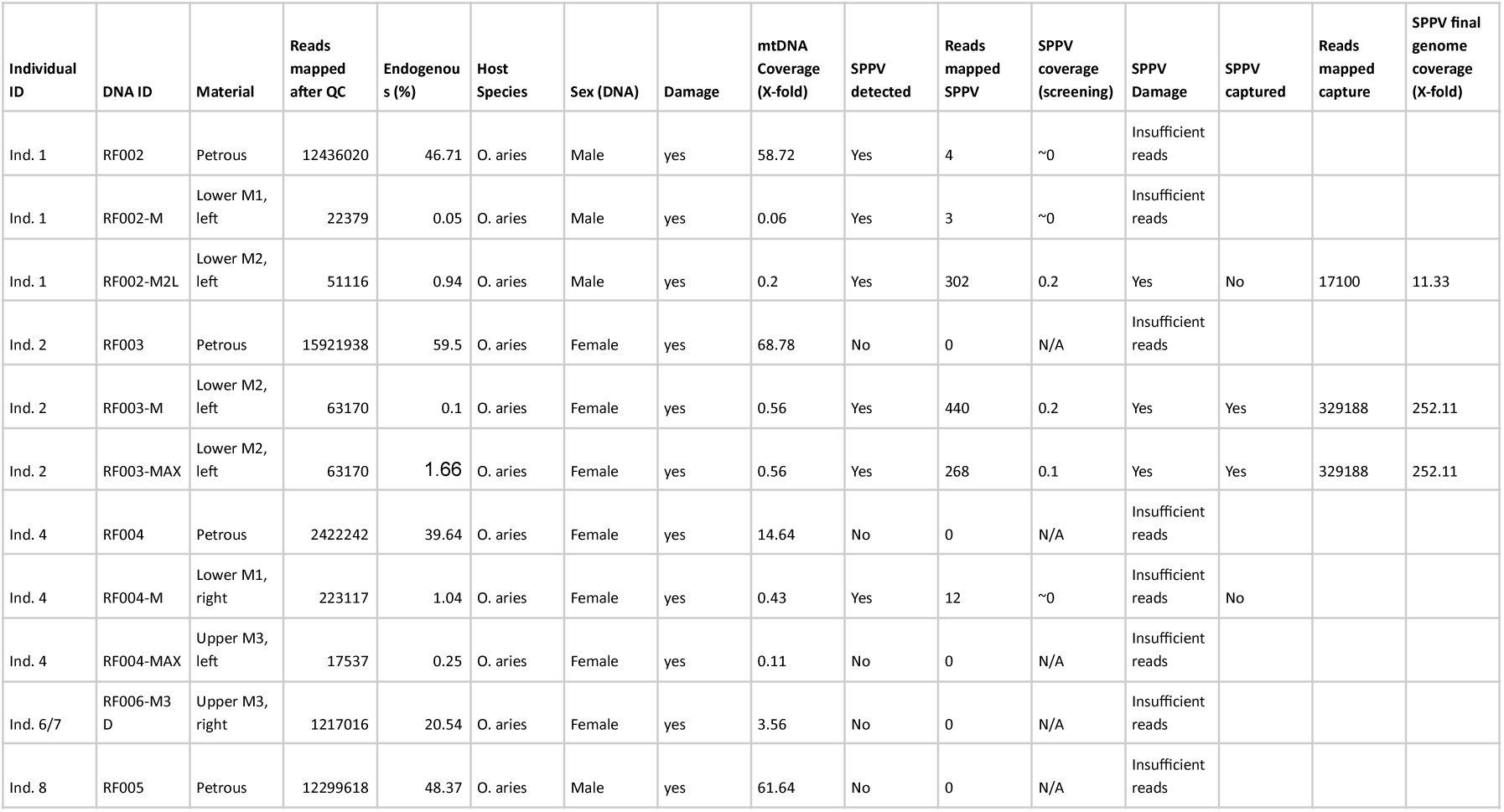
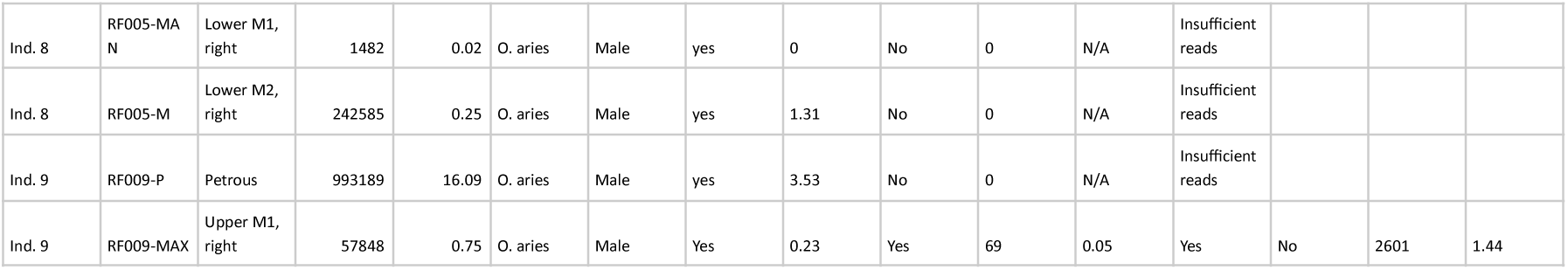
Samples selected for pathogen identification and results of DNA analysis.

| Individual ID | DNA ID | Material | Reads mapped after QC | Endogenous (%) | Host Species | Sex (DNA) | Damage | mtDNA Coverage (X-fold) | SPPV detected | Reads mapped SPPV | SPPV coverage (screening) | SPPV Damage | SPPV captured | Reads mapped capture | SPPV final genome coverage (X-fold) |
| --- | --- | --- | --- | --- | --- | --- | --- | --- | --- | --- | --- | --- | --- | --- | --- |
| Ind. 1 | RF002 | Petrous | 12436020 | 46.71 | O. aries | Male | yes | 58.72 | Yes | 4 | ~0 | Insufficient reads |  |  |  |
| Ind. 1 | RF002-M | Lower M1, left | 22379 | 0.05 | O. aries | Male | yes | 0.06 | Yes | 3 | ~0 | Insufficient reads |  |  |  |
| Ind. 1 | RF002-M2L | Lower M2, left | 51116 | 0.94 | O. aries | Male | yes | 0.2 | Yes | 302 | 0.2 | Yes | No | 17100 | 11.33 |
| Ind. 2 | RF003 | Petrous | 15921938 | 59.5 | O. aries | Female | yes | 68.78 | No | 0 | N/A | Insufficient reads |  |  |  |
| Ind. 2 | RF003-M | Lower M2, left | 63170 | 0.1 | O. aries | Female | yes | 0.56 | Yes | 440 | 0.2 | Yes | Yes | 329188 | 252.11 |
| Ind. 2 | RF003-MAX | Lower M2, left | 63170 | 1.66 | O. aries | Female | yes | 0.56 | Yes | 268 | 0.1 | Yes | Yes | 329188 | 252.11 |
| Ind. 4 | RF004 | Petrous | 2422242 | 39.64 | O. aries | Female | yes | 14.64 | No | 0 | N/A | Insufficient reads |  |  |  |
| Ind. 4 | RF004-M | Lower M1, right | 223117 | 1.04 | O. aries | Female | yes | 0.43 | Yes | 12 | ~0 | Insufficient reads | No |  |  |
| Ind. 4 | RF004-MAX | Upper M3, left | 17537 | 0.25 | O. aries | Female | yes | 0.11 | No | 0 | N/A | Insufficient reads |  |  |  |
| Ind. 6/7 | RF006-M3 D | Upper M3, right | 1217016 | 20.54 | O. aries | Female | yes | 3.56 | No | 0 | N/A | Insufficient reads |  |  |  |
| Ind. 8 | RF005 | Petrous | 12299618 | 48.37 | O. aries | Male | yes | 61.64 | No | 0 | N/A | Insufficient reads |  |  |  |

|  |  |  |  |  |  |  |  |  |  |  |  |  |  |  |  |
| --- | --- | --- | --- | --- | --- | --- | --- | --- | --- | --- | --- | --- | --- | --- | --- |
| Ind. 8 | RF005-MA<br>N | Lower M1,<br>right | 1482 | 0.02 | O. aries | Male | yes | 0 | No | 0 | N/A | Insufficient<br>reads |  |  |  |
| Ind. 8 | RF005-M | Lower M2,<br>right | 242585 | 0.25 | O. aries | Male | yes | 1.31 | No | 0 | N/A | Insufficient<br>reads |  |  |  |
| Ind. 9 | RF009-P | Petrous | 993189 | 16.09 | O. aries | Male | yes | 3.53 | No | 0 | N/A | Insufficient<br>reads |  |  |  |
| Ind. 9 | RF009-MAX | Upper M1,<br>right | 57848 | 0.75 | O. aries | Male | Yes | 0.23 | Yes | 69 | 0.05 | Yes | No | 2601 | 1.44 |

As previously, unaligned reads were then examined for pathogen DNA sequences using *k*-mer matching [20]; whereas the petrous samples yielded no positive results, four teeth samples yielded varying amounts of reads for the sheeppox virus (SPPV, *poxiviridae*): 440 reads (sheep #2), 302 reads (sheep #1), 69 reads (sheep #9) and 12 reads (sheep #4). Alignment-based assessment of this screening data [44] suggested the unambiguous presence of SPPV reads (Figures S4-S7). We additionally observed 3 reads matching SPPV in dental calculus from sheep #2 (Table S3), suggesting it is possible to detect this pathogen in calcified plaque from infected individuals.

Among the four SPPV-positive samples, sheep #2, which showed the highest number of reads aligning to SPPV, was selected for targeted genome enrichment (see Methods), and yielded excellent coverage (252.11×). The SPPV-positive teeth from sheep #1 and #9 and were subsequently re-sequenced after applying USER treatment to remove aDNA-associated damage, resulting in good quality genome coverage for sheep #1 (11.33×) and moderate quality coverage for sheep #9 (1.44×). The recovery of high depth SPPV genomes (range: 1.44–252.11×) from these three individuals strongly implies that each was affected by sheeppox peri-mortem. Sheep #4 yielded too few viral reads from screening shotgun data to enable efficient genome reconstruction from deep sequencing and was therefore not sequenced further. It is likely sheep #4 also was experiencing an active SPPV infection at the time of its death, but we could not recover a complete genome in this instance.

To assess the relatedness between the recovered Bouteiller SPPV genomes and SPPV today, a pairwise distance matrix was computed using single-nucleotide polymorphisms (SNPs), representing positions differing between genomes (Figure 5A). Only two SNPs were observed among the Bouteiller isolates: one unique to sheep #1, and one unique to sheep #2; when we directly examined these loci, both appear to be false positive calls (see Supplementary Note 2 and Figures S8-9), and our recovered SPPV genomes are likely identical. In comparison, 122 SNPs are observed on average between the Bouteiller genomes and modern SPPV, and on average 44 SNPs differentiate pairs of modern SPPV genomes. The recovery of highly similar viral genomes from multiple individuals at the Bouteiller site suggests that the sheep were infected during a single outbreak. Construction of a Maximum Likelihood phylogenetic tree [21] with 16 modern SPPV sequences and 1 Goatpox virus as an outgroup shown that Bouteiller SPPV strains form a distinct, basal monophyletic clade to modern SPPV lineage, consistent with a placement of sequences from the 17th-19th century CE (Figure 5B), as opposed to modern contaminant SPPV genomes.

**Figure 5.**
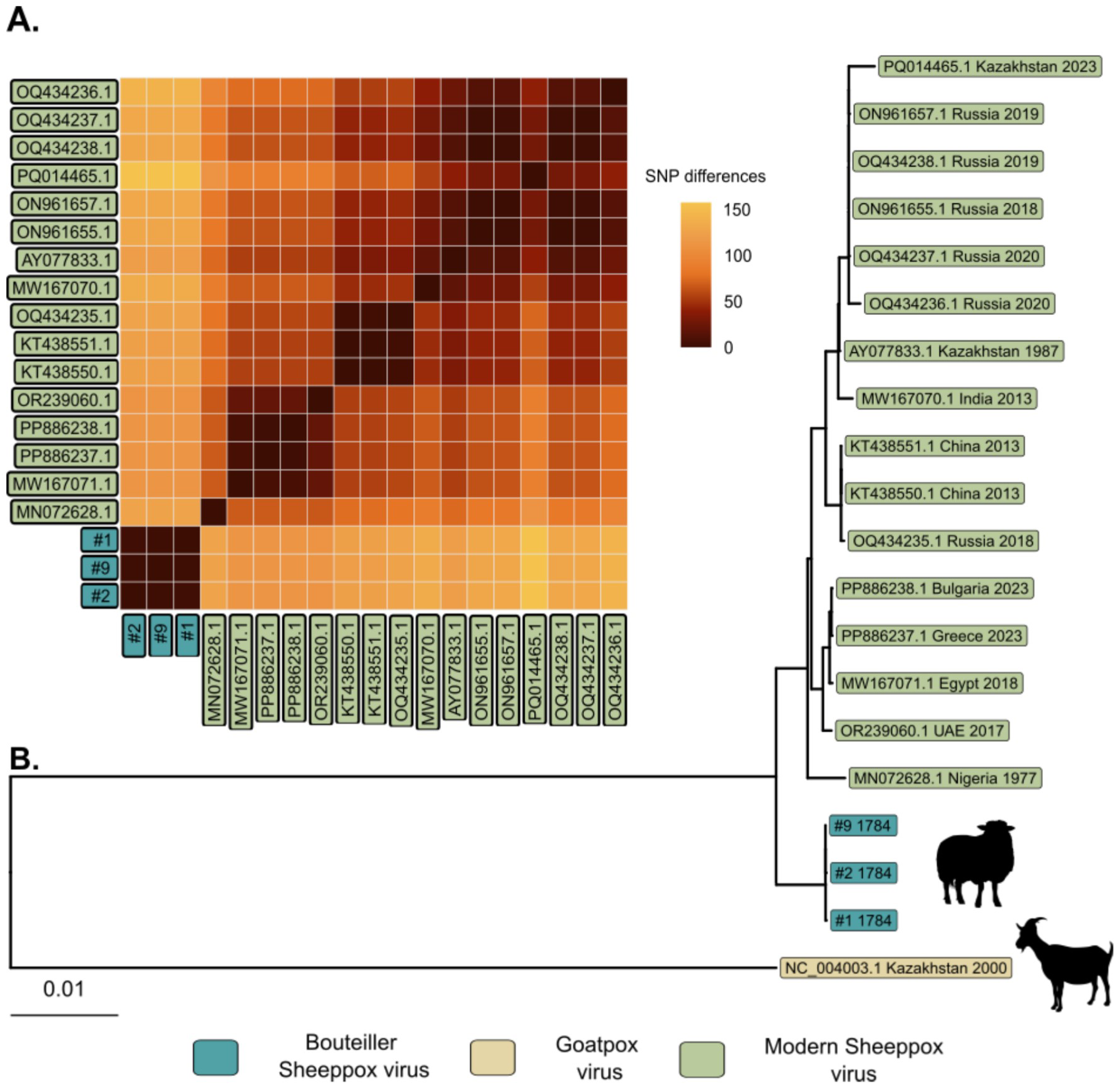
Genomic relatedness and phylogenetic placement of the Bouteiller sheeppox virus isolates. **(A)** Pairwise SNP distance matrix among Sheeppox virus genomes. Colours indicate the number of SNP differences between each pair of sequences, with darker shades representing higher divergence, showing genomic relatedness between the Bouteiller sheeppox virus isolates (#1, #2, #9). Modern isolates are represented by their corresponding GenBank accession numbers matching the isolates from Figure SB. **(B)** Phylogenetic position of the Bouteiller Sheeppox virus isolates relative to modern diversity. A SNP-based Maximum Likelihood phylogeny [21] based on complete genomes, rooted on Goatpox virus. Tip labels show accession numbers and sampling locations and dates. Bouteiller genomes are represented in blue with a sheep silhouette, and the modern Sheeppox virus lineage in green. The outgroup Goatpox virus is represented in beige and with a goat silhouette.

Sheeppox virus is a double-stranded DNA virus of the *Capripoxvirus* genus, within the *Poxviridae* family. It is closely related to goatpox virus (GTPV) and lumpy skin disease virus (LSDV), and more loosely to other members of *Poxviridae*, such as smallpox (variola) or cowpox (vaccinia). Its genome of ca. 150 kbp [45], large for a virus, makes it a good candidate for detection in archaeological material. No report of SPPV identification in archaeological bone has however of yet been published (although *Capripoxvirus* may have been detected from historical manuscripts, see [46]), and this is to the best of our knowledge the first report of a paleogenome of the virus.

## Discussion

### Sheeppox: epidemiology and clinical presentation

Sheeppox is the disease determined in sheep by the SPPV virus. Though other animal species may host the virus, the disease itself is in natural conditions only observed in sheep and very occasionally in goats; cattle appear impervious to most strains [47]. Sheeppox is extremely contagious among sheep, and morbidity in a naïve flock is usually 75% to 100% [48]. It is transmitted through direct contact and infectious aerosols (nasal discharge, saliva, scabs); contagion often occurs during gatherings of animals on communal pastures or at watering points, at fairs and markets, or with the introduction of a new individual in a flock. Mortality is high but variable, with average death rates ranging from 5% to 85% of affected individuals [48,49].

Two main forms of the disease are distinguished. In the regular or benign form, more common in adults, animals initially exhibit fever, loss of appetite, and ocular discharge, followed after four to five days by the eruption of cutaneous papules on the areas of the body not covered in wool, most notably the face. These papules develop into vesicles which subsequently rupture, spreading infectious fluid, and are replaced by nail-like scabs that eventually fall off, leaving permanent scars [50]. Most animals survive, and mortality rates rarely exceed 5% [48]. The malignant form is most frequently observed in animals under two years of age, with which it represents the most common form. The invasion phase is more severe, with marked depression and prostration, and some lambs may die even before the appearance of the papules. The latter affect not only the woolless areas of the skin, but also the mucous membranes and internal organs, and spread to the mouth, lungs, trachea, digestive and urogenital mucosae. Animals may exhibit respiratory difficulties, pneumonia, and hemorrhagic diarrhea, and pregnant females almost invariably abort. In some cases, the eruption may involve the entire skin surface. Mortality rates from malignant sheeppox range from 50-85% of affected animals [48–50]. In survivors of both forms, sheeppox confers a strong and lasting immunity that prevents reinfection.

Terrain has little influence over the prevalence of sheeppox, which mostly depends on local sheep density and the intensity of livestock movement and trade. Its seasonality is debated; the disease seems possible all year round, but various authors point to different preferred seasons, perhaps due to local herding practices and commercial circumstances. Certain breeds are more resistant than others; wool sheep show greater susceptibility than hair sheep, and breeds specifically selected for fine, dense wool such as the merinos are particularly vulnerable. Age is nonetheless the greatest risk factor, with deaths highest in animals aged under 2 years, and more so between 4 and 12-18 months of age: mortality rates can reach 80% in that cohort while most adults of the same flock survive [50]. Among adults, health status is the main risk factor, with aged individuals, those weakened by intercurrent disease and ewes in late pregnancy displaying higher mortality rates [51]. Females appear slightly more at risk than males [50,51].

As sheeppox does not affect the skeletal system, the disease leaves no identifiable traces on archaeological bones. The skin lesions however severely devalue the hides, sometimes rendering them unusable [48], and the repulsive appearance of the lesions may discourage the consumption of the carcasses.

### The history of sheeppox in France

Owing to its characteristic dermatological symptoms and highly contagious nature, sheeppox is easily identified both in live animals and in written accounts, and cannot be mistaken in its typical presentation. Ancient textual evidence can therefore be considered fairly reliable. The earliest written description of the disease dates back to Roman Antiquity, with Columella’s description of the disease of sheep known as “the pustule” ( *pusula*), an extremely contagious and highly lethal dermatological ovine disease (1 ^st^ century CE, [52]). Sheeppox appears to have been fairly common in medieval Western Europe. The disease appears in husbandry treatises such as Walter of Henley (13 ^th^ c. England, [53]) and Jean de Brie (14^th^ c. France, [54]), and is recorded as an explanation for sheep losses in accounting documents all through the 13 ^th^ to 15^th^ centuries in France and England [16]. The contagiousness and lethality of sheeppox were well recognized at the time, and as early as 1499, an ordinance of the *Mesta general de Castilla y León* (Spain) required shepherds to declare all places in which the disease had been observed [55].

In France, mentions remain common throughout the 15 ^th^ and 16^th^ centuries, but appear to dip temporarily in the 17^th^ century, possibly affording flocks a short respite [16]. The disease reappears however with a vengeance in the mid-18 ^th^ century, with epizootic outbreaks increasingly severe in both victim count and geographic spread attracting growing attention from the developing field of veterinary science. At least three treatises are specifically devoted to the disease during that period, as well as numerous chapters and papers [55–59]. In the same period, sheeppox also became the subject of public health policies, with rulings and decrees from 1714, 1778, and 1784 making it a notifiable disease and regulating sheep movement in areas of infection [60]. By the end of the 18th century, the disease had become so common that Claude Bourgelat, founder of the first veterinary school in 1761, “claimed that no animal ever reached the end of its life without having experienced sheep pox.” [61].

In 1801, an ordinance from the Paris Police Prefecture was passed in response to an outbreak affecting the western and northern parts of the region, including Saint-Denis, a major livestock trade centre less than 20 kilometres from Louvres [60]. In addition to organizing the isolation of the sick and prohibiting their sale, the text also prescribes how the victims’ carcasses should be managed: they “shall be buried on the same day with their skin and wool, at a depth of 1.34 meters (4 feet), outside the boundaries of the municipalities, all at the owners’ expense”. Steep fines awaited offenders.

Following the success of smallpox inoculation, experiments with sheeppox inoculation were attempted in the 18^th^ century [62]; though mostly effective, the method never really caught on beyond scientific circles. In the late 19 ^th^ century, the development of an effective and reliable vaccine was met with public success, and combined with stringent public health measures, enabled the eradication of sheeppox from mainland France at the beginning of the 20 ^th^ century [55].

### Interpretation and discussion

In light of this information, the diagnosis of sheeppox obtained by paleogenomic analysis appears consistent with the archaeological data from the Louvres assemblage. The disease is both highly contagious and deadly in sheep, and could easily lead to the simultaneous deaths of nine individuals. Although SPPV infection was only confirmed in three animals, their situation within the pit (the two top-most and the bottom-most individuals), the archaeological evidence of a single burial event followed by quick infilling of the pit, the genomic evidence of a single viral strain, and the extreme contagiousness of sheeppox in naïve animals together point to all nine animals having been affected by the disease. The lack of evidence from the three other animals that were sampled is likely due to differential preservation of the genetic material rather than absence of infection – it is indeed noteworthy that the viral material was only identified in one of the two teeth sampled from individual 1, and in no petrous bones including those from animals which yield SPPV-positive teeth (Table 2).

The deposit also stems from an area and a period in which sheeppox was common: in addition to the 1801 outbreak referenced above, Delafond notes outbreaks in the Parisian area in 1773-1774, 1776, 1805 and 1810 [60]. Hurtrel d’Arboval further states that “In most of the departments of France, [the disease] returns epizootically only once in ten or fifteen years; but its recurrence is more frequent in the vicinity of Paris, where there is much passage, and where a considerable trade in sheep is carried on.” [61]. This reported frequency prevents us from ascribing the Louvres mortality to a specific epizootic event.

If place and time are therefore very typical, the same cannot be said for the demography of the victims. It is established juveniles are most susceptible to sheeppox, and the majority of deaths are usually found within the 4-18 months cohort [50]. However, the youngest animal from Louvres was a subadult around three years old, and the remaining eight animals are mature adults, several of which appear in their senior years. The population is therefore much older than expected, which explains the bad fit of the paleo-epidemiological approach. No definitive explanation can be given for this discrepancy, but several factors might have played a part. First of all, the initial composition of the flock is unknown – perhaps did the farmer not own any lambs or yearlings, or did he keep these animals on another site. The composition of the death assemblage is similarly uncertain: only 130 square meters of the farmyard were excavated during the assessment, and nothing precludes there having been one or several other sheep-pits on the site, which may have included youngsters. This is unfortunately unverifiable, as the site now hosts a large residential building.

Secondly, the Bouteiller sheep do fit the typical profile for adult victims of sheeppox. The latter are indeed most often individuals weakened by age or disease, which is very much in agreement with our results: at least four of the nine victims are identified as older adults above six years of age, and one, sheep 2, displayed strong evidence of *Taenia hydatigena* parasitism. High parasitic burdens are known to contribute to sheeppox mortality [50,56]. Though no information relating to *T. hydatigena* specifically is available, Hurtrel d’Arboval emphasises the severity of the disease in cases of prior liver fluke (*Fasciola hepatica)* infestations [56], the symptoms of which *T. hydatigena* is reported to occasionally mimic [48]. Finally, the high proportion of adult males and the slender morphology of all animals suggest the flock might have been specialised towards the production of wool. Sheeppox is described as being more severe in breeds selected for fine wool, including in the merinos, which was popular around Paris at the time; though no evidence of the specific breed affinities of the Louvres sheep is available, this may have been a contributing factor.

While several hypotheses may account for the discrepancies in age profiles, a temporal evolution in disease presentation can be ruled out. The deaths occurred indeed recently enough that our analysis of sheeppox could rely on multiple, contemporaneous descriptions of its clinical manifestations [14,49,56,57,61].

The case also provides an opportunity to analyse how Modern period societies dealt with contagious disease among their livestock. The earliest legislation relative to sheeppox is dated in France to 1714, with several updates through the late 18 ^th^-early 19^th^ century, which means that unless the deposit is dated from the earlier end of the probability distribution (1658 - 1698 CE (19.0%)), the farmer was probably aware of the existence of regulations –Louvres is not an isolated town. These regulations are possibly what led to the burial of the animals, though it was done only in partial compliance to local ordinances. Whereas the latter prescribe deep burials (4 to 6 French feet, i.e. 1,34m to 2m) in areas outside the town, the Bouteiller sheep were interred directly within the farmyard, less than 60m from the main house, in a pit that was originally only 60 to 80cm deep. At least one animal had been flayed, in violation of all regulations – diseased skins and their trade being a known cause for the spread of the contagion. The diagnosis of sheeppox does allow us to better understand the atypicality of the flay marks on sheep #5: while they appear needlessly high up the limb for regular skinning marks, their location would have allowed cutting off the skin well into the wooled area, avoiding the poxes that would be found on the lower limbs, and therefore avoiding detection as a pocked skin. It is possible that several other individuals from the pit, perhaps all, were similarly flayed, but that a skilled skinner left no marks doing so. All in all, it seems as though the burial reflects only minimal compliance to regulations, either to provide an appearance of compliance, to save time, and to limit losses by selling skins, or out of ignorance of the regulations, perhaps because the burial predated their introduction. Finally, mention should be made of the possibly culled animal. If the traces that are observed are indeed slaughter marks, they could either stem from a mercy killing, intended to shorten the suffering of a dying animal, or from a sanitary precaution, eliminating a diseased sheep that could continue spreading the infection within the flock. Either act is possible within the cultural context, and both are documented in contemporaneous veterinary writings.

## Conclusion

We present a multi-disciplinary approach to investigating the archaeological and biological context of mass morality events in the archaeozoological records. By combining archaeological, archaeozoological, paleopathological, and genetic approaches, we disentangled the health status and likely cause of death of an ovine assemblage from Louvres, France. Extensions of this approach are evident, including the use of isotope analysis and population genetics to confirm local sources of the animals. Our results additionally represent the first genetically confirmed evidence of an ancient animal mass mortality event associated with disease outbreak (sheeppox virus), and recover the first ancient *Taenia* genome. Overall, our approach provides a model for future multi-disciplinary assessments of mass mortality events in the archaeozoological record, thus improving our understanding of animal disease and welfare in the past.

## Methods

### Zooarchaeological methods

The assemblage was analysed according to guidelines for best practices in zooarchaeology [63]; all bones were identified to anatomical part, weighed to a 0.1g precision and measured with digital calipers following the recommendations of Von den Driesch [64]. Species distinction between sheep and goat was carried out using published criteria from [65,66], and sex identification was based on pelvic conformation relying on a combination of criteria from [67–70]. Wither height was assessed by averaging height reconstructions for all long bones from a given individual [71] and slenderness was calculated on the metacarpal using [72]. The sheep’s age-at-death was estimated through tooth eruption and wear for the animals who had preserved teeth [73] and relying on epiphyseal fusion stages for the remaining three individuals (3, 5, 7) [74].

### Paleo-epidemiological methods

In previous work [16], evidence of sheep mass mortalities was collected in French and English medieval and Early Modern written sources, including historical texts (chronicles and annals), economic data (mostly from manorial records), and didactic texts (agricultural and veterinary treatises). All mentions of cause of death were noted, and where relevant, specific diseases were identified on the basis of their symptomatology and designation (see [75]). This allowed us to identify a list of the most common causes for mass ovine mortalities in different time periods; for the 17^th^-19^th^ centuries, disease-related causes included liver fluke, sheeppox, anthrax and Sologne red disease, and accidental causes included grain overload, hunger, hypothermia, wolf and dog predation, lightning strike and drowning [16]. Each of these causes of death was then described according to 16 parameters (historical presence, geography, terrain, seasonality, contagiousness, mortality rates, affected species, racial influence, age profile, sex profile, impact of physiological status, impact of sanitary status, bone lesions, carcass aspect, others), taking into account both historical data describing their past presentation and present-day veterinary knowledge. This information was summarized in twelve fact sheets available in [9].

The parameters obtained from the archaeological and osteological analysis of the Louvres deposit were then compiled into an epidemiological profile (Table S2) and compared to that of the twelve mortality causes, to evaluate fit and identify potential diagnostic hypotheses.

### Ancient DNA laboratory work

All ancient DNA laboratory work was performed in dedicated aDNA facilities in Trinity College Dublin, Ireland. Personal Protective Equipment (PPE) was used at all times, including boiler suits, hair nets, gloves (two layers), and face masks. All equipment and surfaces were regularly cleaned with dilute (0.5%) sodium hypochlorite. PCR amplification reactions were performed in different laboratory facilities to avoid contamination of amplificed materials.

A total of five ovine petrosal bones, ten teeth, and one dental calculus sample were processed. Petrosal bones were subsampled using a Dremel saw at the densest region. Bone fragments were then pulverised using a MixerMill at 30 Hz until fully pulverized. Teeth were sectioned along the cementoenamel junction, and each root was further halved. Material from the pulp chamber and inside root was collected using a dental drill, yielding between 8–59 mg of powder per tooth. Dental calculus from sheep #2 was removed using a sterile blade, generating a total of 20 mg of material.

### Bones, teeth and calculus extraction

Bone, tooth, and calculus material were first subjected to a 15-30-minute pre-digestion in EDTA (15 minutes only for the calculus sample). For teeth samples, both the EDTA wash and the remaining material pellet underwent a 24-hour digestion in EDTA and proteinase K at 37 °C, following the extraction steps in [76]. After incubation, DNA was purified using a modified pH-adjusted PB binding buffer [76].

### Nodule extraction protocol

The pulverized nodule was digested using a similar protocol as described [76], but removing bleach pre-wash. Briefly, 1ml 0.5M EDTA was added to the nodule powder and also an empty tube (extraction blank). Tubes were incubated at 37 °C for 30 min, with constant rotation. Tubes were briefly centrifuged to 13,300 rpm and the EDTA removed and stored. An extraction buffer was prepared (40 μl 1M Tris-HCl, 34 μl SDS, 1880 μl 0.5M EDTA, 26 μl proteinase K), subjecting the buffer to 30 minutes of UV prior to the addition of proteinase K. 1 ml of extraction buffer was then added to each tube and the tubes vortexed. Tubes were incubated for ∼24 hours at 37 °C and gentle agitation.

After digestion, tubes were spun at 13,300 rpm for 10 min to separate supernatant and any remaining nodule. The supernatant was then subject to purification using Large Volume Roche columns and 13 μl of a modified PB buffer (added to 500 ml modified PB: 16 ml 3M sodium acetate, 13.2 ml 5M sodium chloride). After spin down and purification, DNA was eluted in 50 μl EBT.

### Sequencing library building

All libraries were dsDNA libraries constructed following the Meyer and Kircher protocol [77]. One library for each sample was prepared without uracil DNA glycosylase treatment using 12-16 μl starting DNA, to capture native damage patterns. All other prepared libraries were pre-treated with 5 μl UDG-glycosylase (New England Biolabs, M5505L) at 37 °C for 1 hour. Following dual indexed-PCR amplification for 12 cycles, libraries were shotgun sequenced on an Illumina Novaseq X (150 bp pair end). Sequencing statistics and metadata for each indexed amplification are provided in Table S3.

### Sequencing data processing and pathogen detection

Raw sequencing reads were trimmed and quality filtered using AdapterRemoval v2.3.2 [78], discarding reads shorter than 30 bp and merging overlapping pairs (--collapse --minadapteroverlap 1 --adapter1 AGATCGGAAGAGCACACGTCTGAACTCCAGTCAC --adapter2 AGATCGGAAGAGCGTCGTGTAGGGAAAGAGTGT --minlength 30 --trimns --trimqualities). To remove host-derived sequences, libraries were aligned to a concatenated reference comprising human and multiple animal genomes (GRCh38, Sscrofa11.1, ARS-UCD1.2, mCerEla1.1, ASM170441v1, EquCab3.0, ARS-UI_Ramb_v2.0). Metagenomic profiles were generated from the resulting unmapped reads using KrakenUniq v1.0.4 [20] and a custom database built from a microbial version of the NCBI “nt combined” dataset supplemented with human and complete eukaryotic reference genomes.

To evaluate the reliability of taxonomic assignments, we applied the scoring framework of Guellil et al. [79], to compute an E value calculated at the species level to distinguish between true and false positives in KrakenUniq taxonomic assignment. Taxa were subsequently filtered for a predefined panel of animal and human pathogens ( Table S5), retaining only assignments with E value > 0.001 and ≥ 20 supporting reads. To validate the E score signal of Sheeppox virus in the Bouteiller samples, we ran a mapping-based detection with HOPS [44] on positive libraries using the approach and database described in [8].

### RNA bait enrichment for sheeppox virus DNA

A set of custom in-solution 3x-tiled 80bp RNA baits (myBaits®) were designed and synthesized by Daicel Arbor Bioscience, targeting *Capripoxvirus* DNA. The bait design was based on reference grenomes sequences for *Capripoxvirus* members (Sheeppox virus: NC_004002.1; Goatpox virus: NC_004003.1; Lumpy Skin Disease Virus, NC_003027.1), with low complexity regions masked. Baits matching to host livestock genomes (Sheep: GCF_016772045.2; Goat: GCF_001704415.2_ARS1.2; Cattle: GCF_002263795.3_ARS-UCD2.0) were excluded (5 baits), as were baits with more than 35% repeat content. The final set of 80bp baits amount to 15,646 unique probes, designed to enrich across *Capriopoxvirus*.

### Capripoxvirus capture application

Amplified, dual index-incorporated libraries were targeted for enrichment for SPPV DNA, through application of the RNA baits described above. Amplified, multiplexed libraries were combined into a pool. The pool was desiccated and worked back up to 7 μl of lab grade H _2_O. The pool was then subject to overnight (∼20 hours) capture using the RNA baits, following the manufacturer’s instructions and making recommended modifications for degraded DNA (“High Sensitivity”), including setting the the annealing temperature to 55 ॰C and performing a second round of annealing and amplification (8 cycles) after the first amplification (16 cycles). Amplification was performed using KAPA HiFI Hotstart (Roche). Purified, captured libraries were then subject to shotgun sequencing for 150bp paired end reads on NovaSeq X platforms (Macrogen Europe, Amsterdam).

### Analysis of sheep DNA

To analyze host DNA, the collapsed reads were aligned on the sheep reference genome Rambouillet v2 (ARS-UI_Ramb_v2.0) using the bwa aln algorithm [80] with relaxed parameters (-l 1024 -n 0.01 -o 2). Reads with mapping quality under 30 were filtered out with samtools v1.19.2 [81], and duplicates were removed with picard MarkDuplicates v2.26.11 (https://github.com/broadinstitute/picard). We calculated post-mortem damage patterns on the *Ovis aries* alignment using mapDamage2 [82], and are shown in Figure S3.

To determine the genetic sex of sheep individuals, we assessed the ratio of reads aligning to autosomal and X chromosome contigs vs contig length [83]. Individuals with two X chromosomes should show the same read number-contig length relationship among investigated chromosomes, while individuals only carrying one X chromosome (that is, genetic males) should show depressed read alignment for X chromosome contigs. We report the genetic sex of each individual in Table 1 and 2.

To assess allele sharing between sheep genome alignments, we used ANGSD [84] to calculate D statistics among sheep alignments for the nodule (RF001) and four of the petrous bone specimens (RF002-RF005, corresponding to individuals 1, 2, 4 and 8 respectfully). The following parameters were used: -doAbbababa 1 -doCounts 1 -minQ 20 -minMapQ 30 -rmTrans 1. The D scores and bootstrapped standard error (SE) estimates for these comparisons are presented in Table S4.

Consensus mtDNA sequences were called from petrous and nodule aligned data on the sheep mitochondrion (NC_001941.1) with angsd doFasta (-doCounts 1 -trim 4 -setMinDepth 3 -setMaxDepth 100 -minQ 20 -minMapQ 30). A multiple sequence alignment was then produced by aligning consensus sequences with NC_001941.1 by using MAFFT with default parameters. A Maximum Likelihood tree was built with IQ-TREE2 [21] employing the nucleotide substitution model HKY+F as determined by ModelFinder [85].

### Analysis of Taenia hydatigena DNA

To validate the presence of *Taenia hydatigena* in the nodule DNA we aligned host (sheep)-removed reads on the the mitochondrial genome of the *Taenia hydatigena* parasite (NC_012896.1), using the same steps and parameters using for alignment to the sheep genome. A consensus mtDNA sequence from the resulting bam file was then produced by using angsd doFasta (-doCounts 1 -trim 4-setMinDepth 3 -setMaxDepth 100 -minQ 20 -minMapQ 30). To validate the *T. hydatigena* identity of the nodule DNA, the consensus sequence was aligned using MAFFT [86] with *Taenia asiatica* (NC_004826.2), *Taenia crassiceps* (NC_002547.1), *Taenia ovis* (NC_021138.1), *Taenia pisiformis* (NC_013844.1), *Taenia saginata* (NC_009938.1), *Taenia solium* (NC_004022.1) and *T. hydatigena* (NC_012896.1) mitochondrial genome. A Maximum Likelihood phylogeny of *Taenia* diversity was built with IQ-TREE2 [21] employing the nucleotide substitution model TIM+F+I+G4 as determined by ModelFinder [85] and rooted on *T. Crassiceps* (NC_002547.1).

After validating the phylogenetic placement of the Bouteiller nodule DNA as being *T. hydatigena* we decided to test for phylogenetic placement of Bouteiller nodule DNA into *T. hydatigena* modern mitochondrial diversity. The same process was applied as described below by using the full diversity of A and B *T. hydatigena* haplogroups (see Figure 4B). The maximum likelihood was built using the nucleotide substitution model TN+F+R2 as determined by ModelFinder [85] and the tree was midpoint rooted.

### Analysis of sheeppox virus DNA

After alignment on the host, unaligned UDG treated and captured libraries were aligned to a modified Sheeppox virus (SPPV) reference genome (NC_004002.1) using the bwa aln algorithm with relaxed parameters (-l 1024 -n 0.01 -o 2). The SPPV genome contains two identical inverted terminal repeat (ITR) regions at its extremities, causing reads derived from these regions to map ambiguously to both ends and receive a mapping quality of zero, which would remove them from downstream analyses. To avoid this, we truncated the terminal 2,213 bp from the reference genome using seqkit [87], allowing unique alignment of reads to the remaining extremity. Alignments with a mapping quality <30 were discarded using samtools v1.19.2, and PCR duplicates were removed with Picard MarkDuplicates v2.26.11. BAM files originating from the same samples were subsequently merged using Picard MergeSamFiles v2.26.11.

Consensus sequences were generated by calling variants with FreeBayes v1.3.6 (https://github.com/freebayes/freebayes), using minimum coverage thresholds of 100× for individual 1, 6× for individual 2, and 1× for individual 9, and requiring ≥90% support for alternative allele calls (--report-monomorphic --min-alternate-count mincov --min-coverage mincov --min-mapping-quality 30 --min-alternate-fraction 0.90 --ploidy 1). Variant call files were then filtered with bcftools view to mark low-quality positions as missing. Final consensus sequences were produced with bcftools consensus, masking missing sites, low-coverage positions, and low-quality variants as N (-a N -M N).

Consensus sequences were then aligned with 16 modern SPPV sequences ( Table S6) and the Goatpox virus reference sequence (NC_004003.1) with MAFFT [86]( --adjustdirection --maxiterate 1000). SNPs were then extracted from the resulting alignment with snp-sites v2.5.1 by extracting only ACGT positions. A maximum likelihood tree was then builded with IQ-TREE2 [21] using the TVMe+ASC substitution model selected by ModelFinder [85] and was rooted on Goatpox virus.

To check for nucleotide variation between the Bouteiller genomes, a SNP distance matrix was computed on the resulting set of SNP with snp-dists (https://github.com/tseemann/snp-dists).

## Acknowledgements

Thanks are due to François Gentili, INRAP, for entrusting A. B.-R. with the study of the animal assemblage excavated in Louvres; to Anicet Konopka for sharing his photographs of the deposit; to Daniel Bradley for access to laboratory facilities at Trinity College Dublin and his mentorship of K. G.D. and L.L’H.; to Harry Little for assistance with laboratory work; to Valeria Mattiangeli for her advice regarding the sheeppox virus capture.

## Data availability

All raw sequencing data generated here is available on the European Nucleotide Archive, under Project Accession PRJEB103957.

