## Supplementary Information for "Evidence of sheeppox in 17^th^-19^th^ century France: a multi-disciplinary investigation of a sheep mass mortality assemblage"

***Supplementary Note 1. Discussion of the location and differential diagnosis of the nodule***

The pathological nodule was recovered from among the cranial debris of individual #2, whose skull rested on the thorax of sheep #3 and beneath the metatarsals of sheep #1. As the distal limbs are rarely prone to ectopic calcification, and any calcification of thoracic origin would have fallen into the void left by decomposition rather than migrated upward through the stratigraphy, it appears likely that if the specimen indeed originated from that area of the pit, it could be attributed to the cranial or cervical region of individual #2.

In animals, the most common forms of cranial or cervical ectopic calcifications are salivary gland calculi (sialoliths) and calcifications of the cervical lymph nodes. The first possibility can be dismissed here, as these calculi are solid and dense, often composed of concentric layers with a rough outer surface [1]. The second possibility corresponds closely to our specimen. Lymph-node calcifications form through hydroxyapatite deposition within and around the lymph node, appearing radiographically as small masses with well-defined yet often irregular peripheries, variable radiopacity—most often heterogeneous—and no clearly defined internal structures [2–5]. Such calcifications are non-specific and develop following chronic inflammation of a lymph node.

Their frequency in ruminants is unknown, as they are most often identified incidentally at slaughter; the calcification itself is asymptomatic. The most frequent underlying causes in sheep are caseous lymphadenitis (*Corynebacterium pseudotuberculosis*) and, less often, tuberculosis (*Mycobacterium tuberculosis*) [6–8]. Cases linked to brucellosis (*Brucella* sp.) or cancer (primary or metastatic) have been recorded in humans and may therefore also be possible in sheep [3,9].

***Supplementary Note 2. Inspection of variant sites called between Bouteiller SPPV sequences***

To validate whether the SNP differences identified between the three Bouteiller Sheeppox virus genomes represented true biological variants, we evaluated the supporting evidence at each polymorphic site (two, covered in all genomes), representing sites 42,598 and 108,583 in the Sheeppox virus reference (NC_004002.1). Variant calls produced by Freebayes during consensus genome generation were first filtered to retain only SNP sites shared between samples, and SNPs differences were extracted and manually inspected using the Integrative Genome Viewer [10].

Both sites are presented in Figures S8 and S9 respectively. For position 42,598, (Figure S8) the variant site overlaps a "GAGG[A homopolymer]" motif. RF003/Sheep #2 shows reads which are called as having an A insertion (bold) and an A variant two bases downstream, leading to a "G**A**AG[A homopolymer]" run at this position. Other reads for this individual, and other SPPV genomes recover,carry no insertion at this position but instead are called as having a different G->A substitution, again leading to "GAAG[A homopolymer]". We conclude the homopolymer stretch is leading to a miscalling of reads at this site for RF003/Sheep #2, and the most parsimonious variant is a single substitution (rather than insertion plus substitution), seen in the other two recovered SPPV genomes. Therefore, the Bouteiller genomes likely do not differ at this site.

Inspecting site 108,583 showed the position to be consistent in all three samples (Figure S9). Freebayes appears to have called this site as varying in RF009/Sheep #9, carrying a "TTTT" variant, while the other two samples carry a "TTTC" variant:

| CHROM | POS | REF | ALT | RF002 | RF003 | RF009 |
|---|---|---|---|---|---|---|
| NC_004002.1 | 108583 | CTTT | TTTC,TTTT | 1 | 1 | 2 |

However, the alignment Figure S9 indicates that all reads across all three samples carry the "TTTC". RF009/Sheep #9 appears to have a single read carrying a "T" variant after the C (i.e. "TTTCT"), which could perhaps affect calling. We concluded that the reads at this site supported all three SPPV genomes carrying the "TTTC" variant, relative to the reference.

*Supplementary Figures*

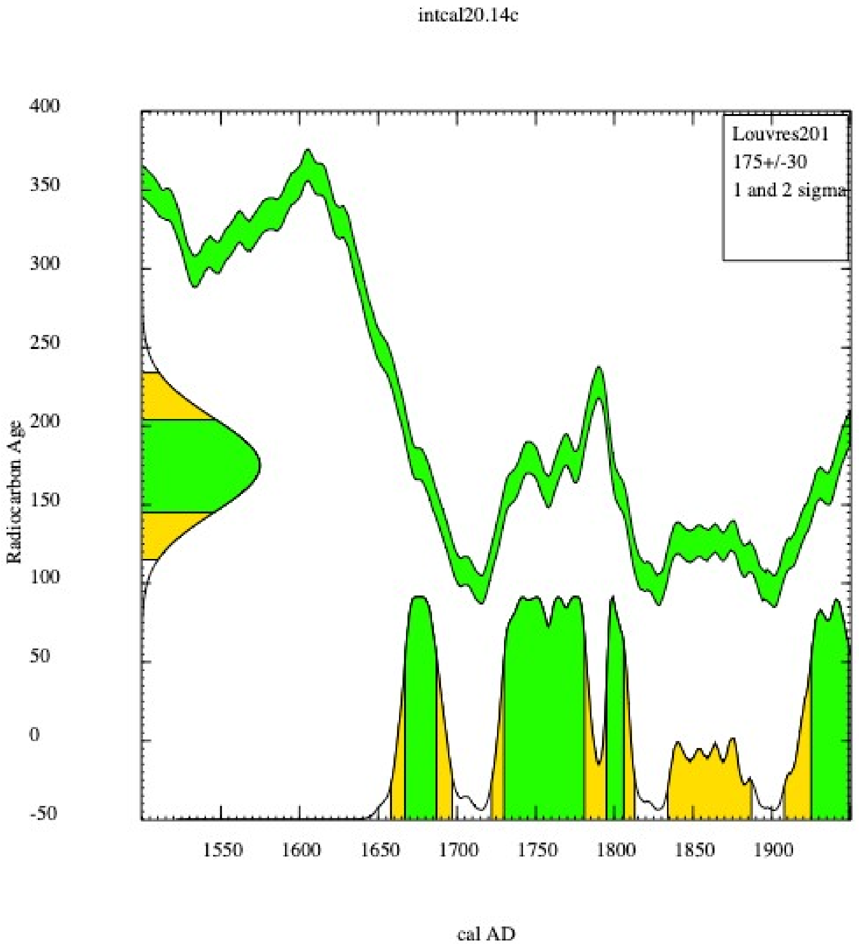


**Figure S1: Radiocarbon age calibration for Louvres 'Le Bouteiller', ind. 2: probability distribution curve. Calibration data set: intcal20.14c, from Reimer et al. 2020.**

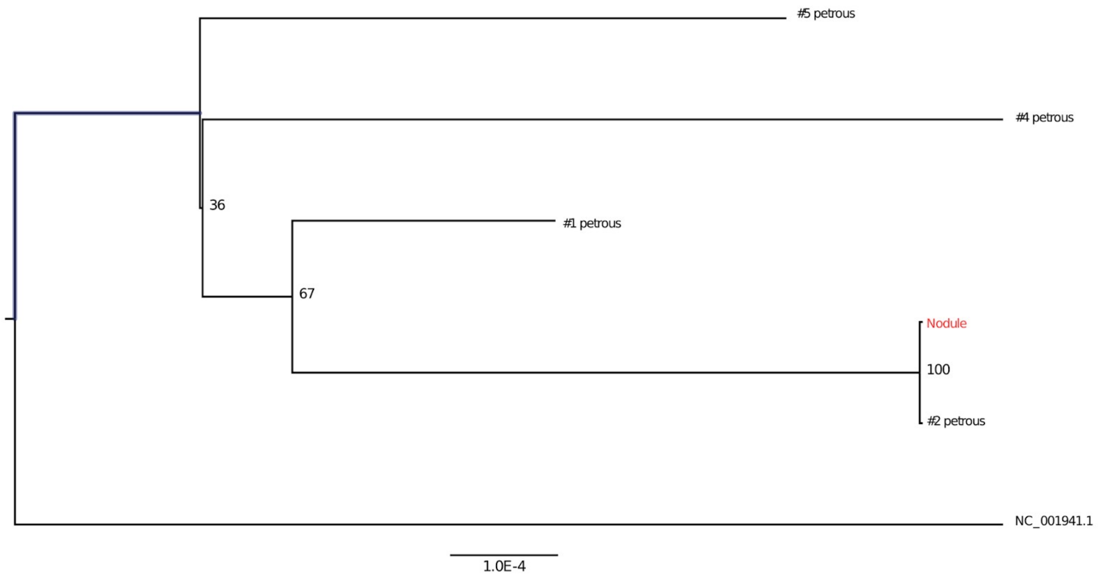


**Figure S2: Maximum likelihood tree of sheep mitochondrial DNA extracted from petrous bones and nodule samples from different individuals of Bouteiller site.** Node bootstrap support values are indicated. The sheep reference mitochondrial DNA reference was included in the alignment and tree (NC_001941.1).

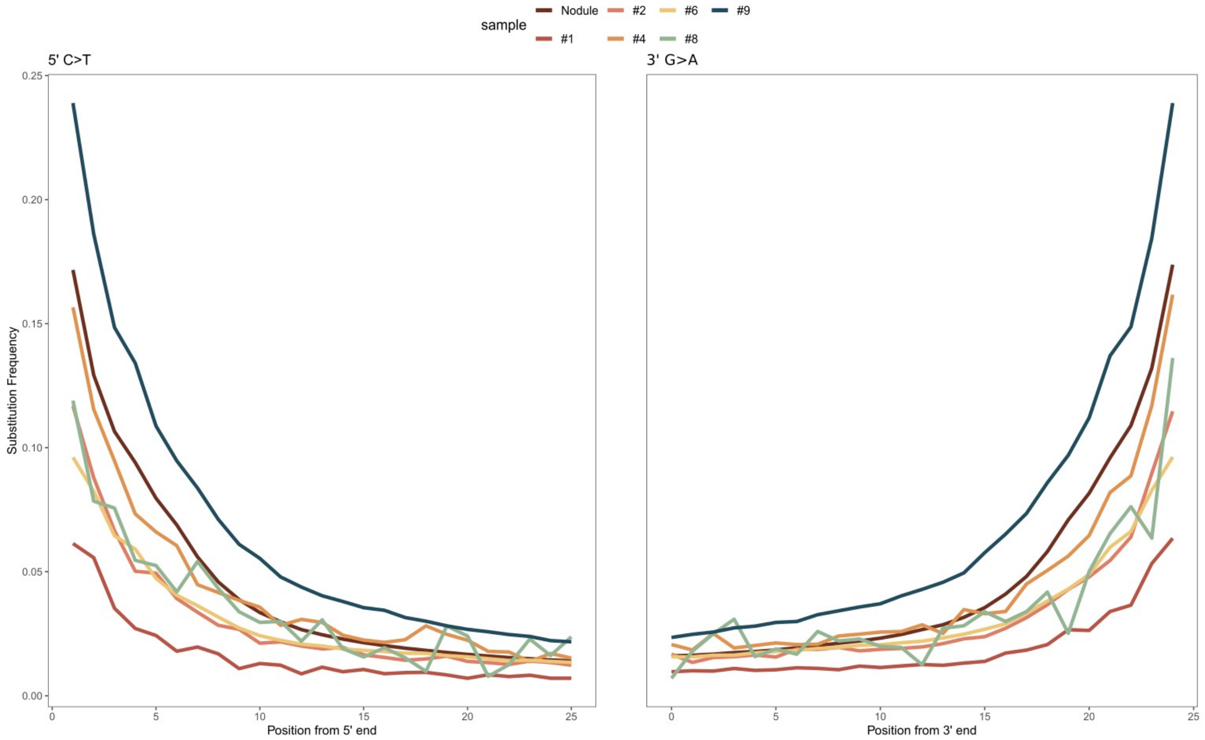


**Figure S3: mapDamage plots of sheep-aligned DNA for Bouteiller sheep and nodule samples, non-UDG libraries.**

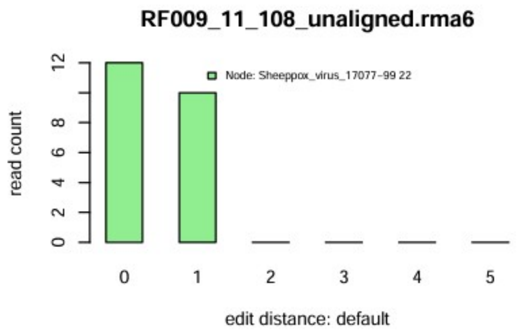


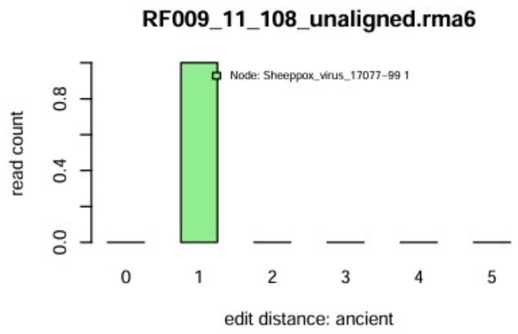


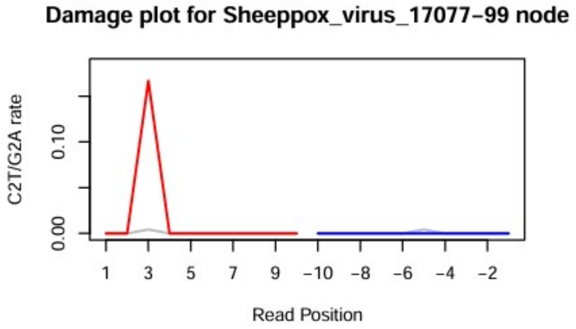


| | |
|---|---|
| Node | Sheeppox_virus_17077-99 |
| Top Reference | Sheeppox_virus_17077-99 |
| all reads | 23 |
| nonDup | 23 |
| readDis | 1 |
| nonStacked | 23 |
| destacking? | On |
| C>T_1 | 0 |
| G>A_-1 | 0 |
| mean length (sd) | 111 (45.943) |

| |
|---|
| Sheeppox_virus_17077-99;_NC_004002.1 100% |
| NA% |
| NA% |
| NA% |
| NA% |
| NA% |
| NA% |
| NA% |
| NA% |
| NA% |

**Figure S4: HOPS pipeline output for RF009 (sheep #9) screening data, with 23 reads assigned to the Sheeppox virus node.**

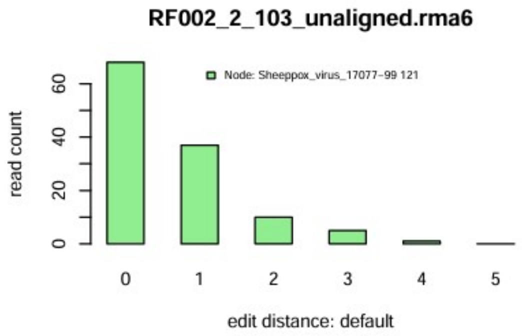


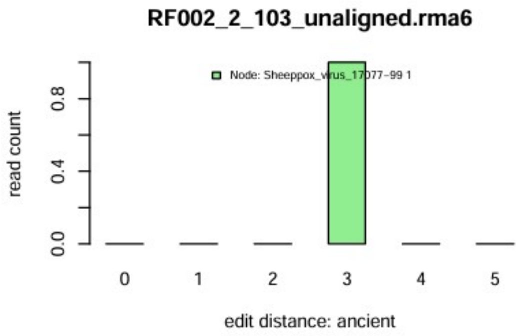


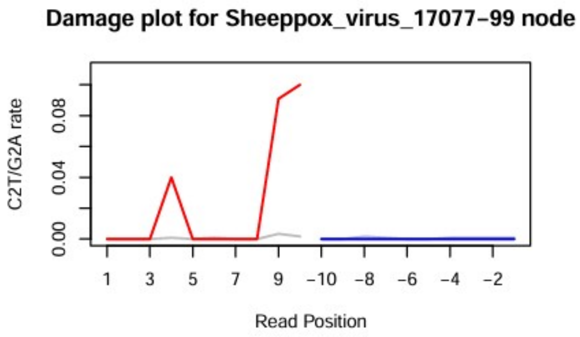


| | |
|---|---|
| Node | Sheeppox_virus_17077−99 |
| Top Reference | Sheeppox_virus_17077−99 |
| all reads | 176 |
| nonDup | 176 |
| readDis | 0.817 |
| nonStacked | 121 |
| destacking? | On |
| C>T_1 | 0 |
| G>A_-1 | 0 |
| mean length (sd) | 113 (37.478) |

| Sheeppox_virus_17077−99;_NC_004002.1 100% |
|---|
| NA% |
| NA% |
| NA% |
| NA% |
| NA% |
| NA% |
| NA% |
| NA% |
| NA% |

**Figure S5: HOPS pipeline output for RF002 (sheep #1) screening data, with 176 reads assigned to the Sheeppox virus node.**

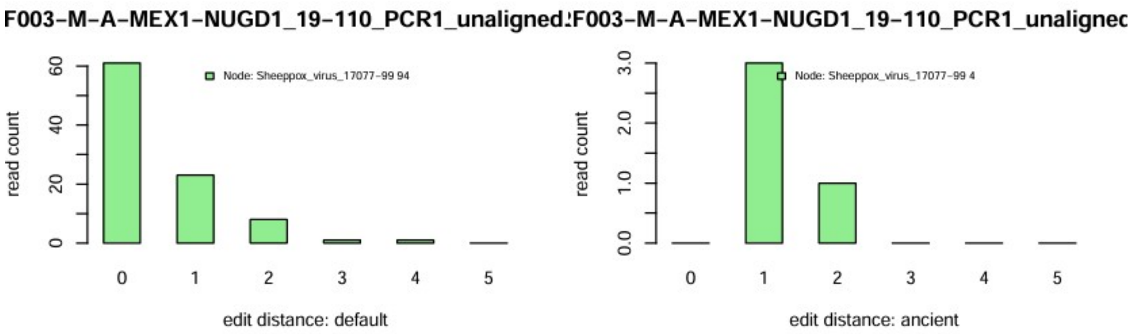


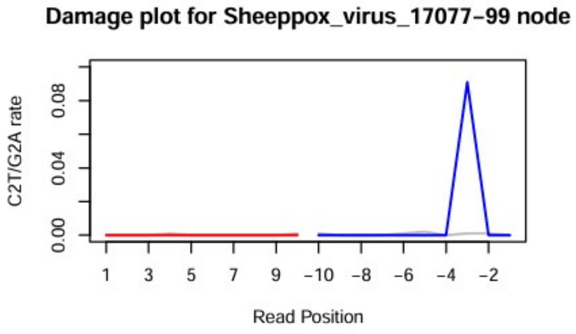


| | |
|---|---|
| Node | Sheeppox_virus_17077-99 |
| Top Reference | Sheeppox_virus_17077-99 |
| all reads | 135 |
| nonDup | 135 |
| readDis | 0.803 |
| nonStacked | 96 |
| destacking? | On |
| C>T_1 | 0 |
| G>A_-1 | 0 |
| mean length (sd) | 109 (42.547) |

| Sheeppox_virus_17077-99;_NC_004002.1 100% |
|---|
| NA% |
| NA% |
| NA% |
| NA% |
| NA% |
| NA% |
| NA% |
| NA% |
| NA% |

**Figure S6: HOPS pipeline output for RF003 (sheep #2) screening data, with 135 reads assigned to the Sheeppox virus node.**

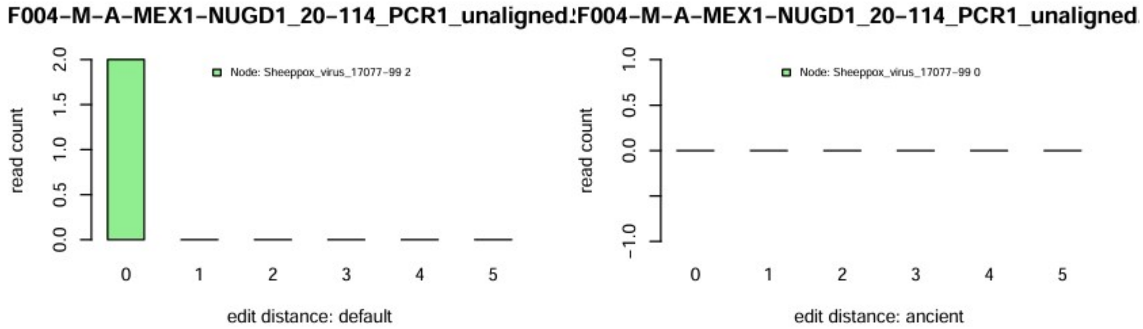


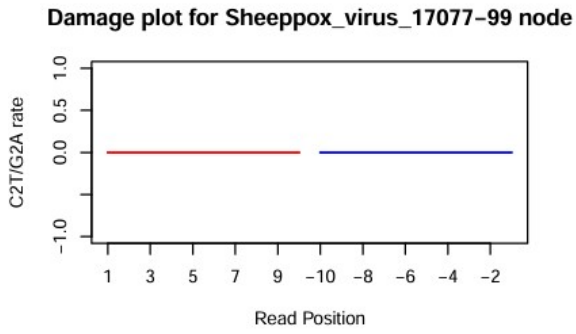


| | |
|---|---|
| Node | Sheeppox_virus_17077-99 |
| Top Reference | Sheeppox_virus_17077-99 |
| all reads | 2 |
| nonDup | 2 |
| readDis | 1 |
| nonStacked | 2 |
| destacking? | On |
| C>T_1 | 0 |
| G>A_-1 | 0 |
| mean length (sd) | 54 (0) |

| Sheeppox_virus_17077-99;_NC_004002.1 100% |
|---|
| NA% |
| NA% |
| NA% |
| NA% |
| NA% |
| NA% |
| NA% |
| NA% |
| NA% |

**Figure S7: HOPS pipeline output for RF004 (sheep #4) screening data, with 2 reads assigned to the Sheeppox virus node.**

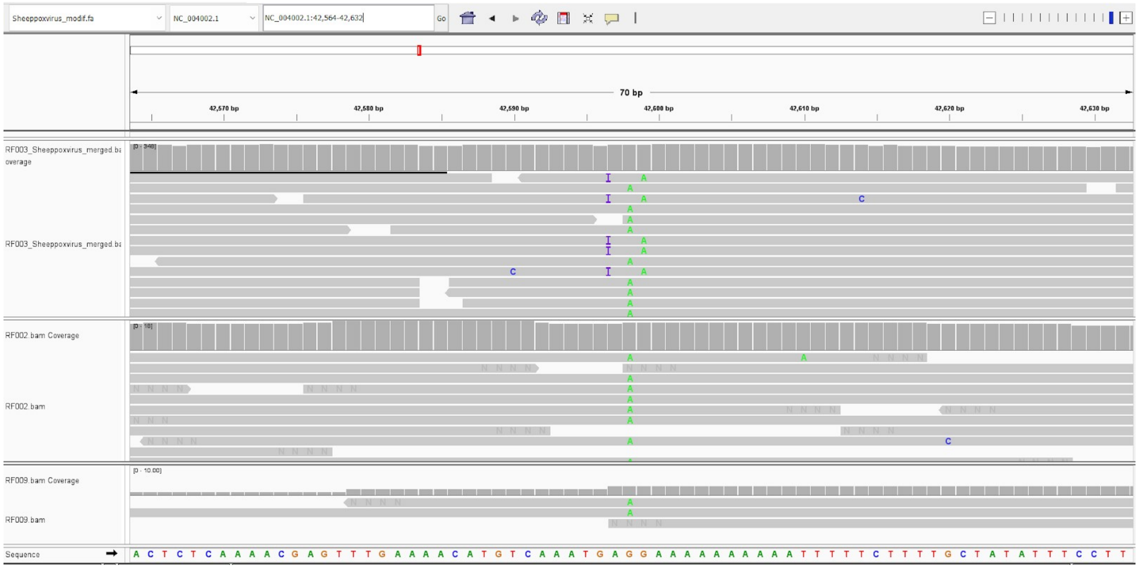


**Figure S8: Visual assessment of Bouteiller SPPV variant site 42,598.**

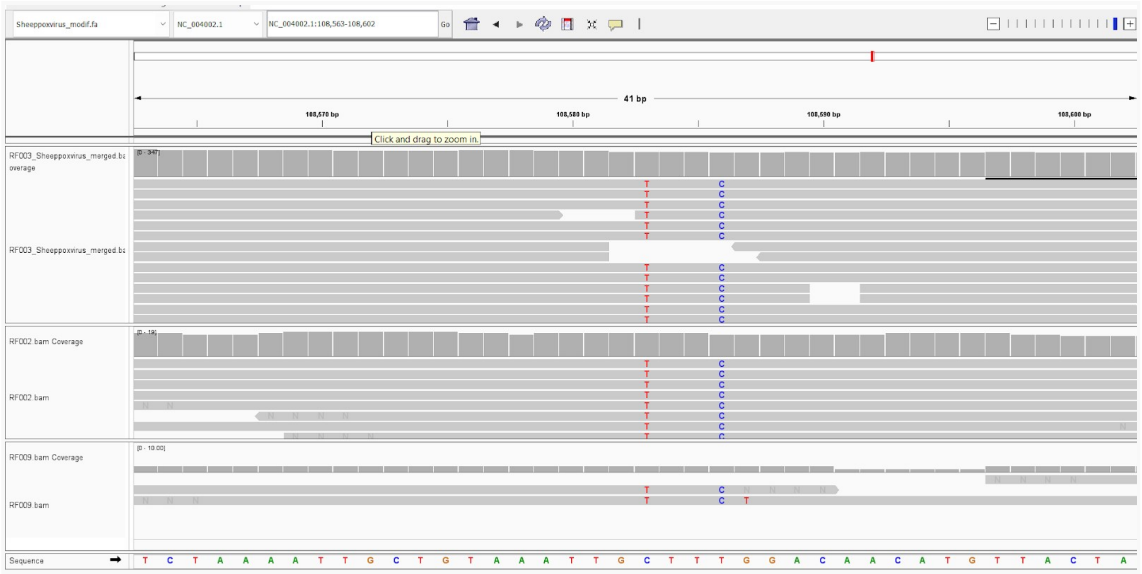


**Figure S9: Visual assessment of Bouteiller SPPV variant site 108,583.**

### *Supplementary References*


1. Polet C, Louryan S, Werquin J-P. 2004 A Mesolithic case of odontoma? *Arch. Oral Biol.* **49 (6)**, 501–506. (doi:10.1016/j.archoralbio.2004.01.008)

2. Gross B, Schneider H, Proto A. 1980 Eggshell calcification of lymph nodes: an update. *Am. J. Roentgenol.* **135**, 1265–1268. (doi:10.2214/ajr.135.6.1265)

3. Tortorich J, Anderson KM. In press. Diagnostic Considerations of Calcified Lymph Nodes. *J. Tenn. Dent. Assoc.*

4. Baud C-A, Kramar C. 1990 Les calcifications biologiques en archéologie. *Bull. Mém. Société Anthropol. Paris* **2**, 163–169. (doi:10.3406/bmsap.1990.1755)

5. Komar D, Buikstra JE. 2003 Differential diagnosis of a prehistoric biological object from the Koster (Illinois) site. *Int. J. Osteoarchaeol.* **13**, 157–164. (doi:10.1002/oa.670)

6. Brugère-Picoux J. 2004 *Maladies des moutons*. Paris: Editions France Agricole.

7. Guimarães A de S, Ribeiro D, Lage AP, Heinemann MB, Miyoshi A, Gouveia AMG. In press. CASEOUS LYMPHADENITIS: EPIDEMIOLOGY, DIAGNOSIS, AND CONTROL.

8. Fontaine MC, Baird GJ. 2008 Caseous lymphadenitis. *Small Rumin. Res.* **76**, 42–48. (doi:10.1016/j.smallrumres.2007.12.025)

9. Eisenkraft BL, Som PM. 1999 The spectrum of benign and malignant etiologies of cervical node calcification. *Am. J. Roentgenol.* **172**, 1433–1437. (doi:10.2214/ajr.172.5.10227533)

10. Robinson JT, Thorvaldsdóttir H, Winckler W, Guttman M, Lander ES, Getz G, Mesirov JP. 2011 Integrative genomics viewer. *Nat. Biotechnol.* **29**, 24–26. (doi:10.1038/nbt.1754)
